# Ablation of the evolutionarily acquired functions of the *Atp1b4* gene in mice protects against obesity and increases metabolic capacity

**DOI:** 10.1101/2025.02.26.640446

**Authors:** Nikolai N. Modyanov, Lucia Russo, Sumona L. Ghosh, Tamara Castaneda, Himangi G. Marathe, Raymond E. Bourey, Sonia M. Najjar, Ivana L. de la Serna

## Abstract

The co-option of vertebrate orthologous *ATP1B4* genes in placental mammals has radically altered the properties of the encoded BetaM proteins, which are genuine β-subunits of Na,K-ATPases in lower vertebrates. Eutherian BetaM acquired an extended Glu-rich N-terminal domain resulting in complete loss of its ancestral function and became skeletal and cardiac muscle-specific component of the inner nuclear membrane. BetaM is expressed at the highest level during perinatal development and is implicated in gene regulation (Pestov et al., Proc Natl Acad Sci U S A. 2007). Here we report the long-term consequences of the *Atp1b4* ablation on metabolic parameters in adult mice. BetaM deficient (*Atp1b4-/Y)* mice have significantly lower body weight and remarkably low adiposity. They exhibit lower fasting blood glucose, enhanced insulin sensitivity, and improved glucose tolerance as compared to their wild type littermates. Knockout mice display higher heat production, increased food intake, elevated oxygen consumption especially in darkness, and higher locomotor activity. The lower respiratory exchange ratio of knockout mice indicates that fat from the diet is metabolized rather than deposited as storage. These robust changes in mouse metabolic parameters induced by *Atp1b4* disruption clearly demonstrate that eutherian BetaM plays an important role in the regulation of adult mouse metabolism. Ablation of *Atp1b4*, leading to the loss of evolutionarily acquired BetaM functions, serves as a model for a potential alternative pathway in mammalian evolution. Essentially, *Atp1b4* ablation simulates a scenario where a specific stage in mammalian evolution is bypassed. Our results suggest that bypassing the co-option of Atp1b4 potentially reduces susceptibility to obesity.

## INTRODUCTION

ATP1B4 genes are present in a single unambiguous copy in all known vertebrate genomes in a highly conserved chromosomal segment located on Xq24 in the human genome (1). The evolutionary history of ATP1B4 genes is rather unique and represents an instance of orthologous vertebrate gene co-option (2) that radically altered properties of the encoded BetaM proteins (1). These proteins in lower vertebrates are authentic Na,K-ATPase β-subunits which in complex with α-subunits function as ion pumps in the plasma membrane (1). In placental mammals, BetaM proteins gained entirely different properties through structural changes of N-terminal domains by acquisition of N-terminal Arg-rich nonapeptide, a typical nuclear localization signal (3), and two extended Glu-rich clusters (1, 4–7). These homopolymeric amino acid repeats typically form intrinsically disordered domains that function as flexible molecular recognition elements in numerous signaling proteins and transcription regulators, enabling interactions with a diverse range of binding partners (8, 9). As a result of these evolutionary modifications, eutherian BetaM has entirely lost its ancestral function as a Na,K-ATPase subunit and has instead become a muscle-specific resident of the inner nuclear membrane, with its acquired N-terminal domain exposed to the nucleoplasm (1, 6, 7, 10–13). Moreover, in sharp contrast to other structurally related members of the Na,K-ATPase β-subunit family, the polypeptide chain of eutherian BetaM is extremely unstable and highly sensitive to degradation by endogenous proteases (10,12). It rapidly disappears in primary culture and is absent in muscle cell lines (7,10,12). BetaM expression is tightly regulated during development, reaching its highest levels in perinatal skeletal myocytes. (6, 7, 10, 11).

We originally reported that in neonatal skeletal muscle, BetaM directly interacts with nuclear transcriptional co-regulator SKIP and is involved in regulation of gene expression as exemplified by augmentation of expression of Smad7, which is negative regulator of TGF-β/Smad signaling pathway (1). We have also shown that expression of exogenous BetaM in cultured muscle cells is sufficient to up-regulate expression of endogenous MyoD, a master regulator of myogenesis, through promoting changes in chromatin structure and enhancing recruitment of SWI/SNF chromatin remodeling enzymes to the DRR region of the MyoD promoter (14).

All these new structural and functional properties acquired through ATP1B4 gene co-option and a unique pattern of expression strongly suggest that eutherian-specific BetaM functions are essential, even might be necessary for survival of placental mammals in natural conditions, and that they provide evolutionary advantage. Moreover, disruption of the ATP1B4 gene may cause long-term physiological and metabolic consequences.

To analyze BetaM function directly *in vivo,* the Atp1b4 knockout mouse model was generated. Initial characterization of *Atp1b4* knockout mice revealed that homozygous *Atp1b4-/-* females have reproductive defects. However, heterozygous *Atp1b4+/-* females and *Atp1b4-/Y* hemizygous males are fertile and generate a progeny that allow direct comparison of wild type and knockout male littermates. Importantly, we have shown that loss of BetaM results in significantly lower body size and weight, growth retardation, and high mortality of *Atp1b4* knockout neonates. These results indicate that BetaM plays an important role during a critical period of perinatal and neonatal development of placental mammals. However, analysis of the long-term metabolic consequences of the *Atp1b4* ablation in adult male littermates revealed a somewhat unexpected and profound beneficial effect of *Atp1b4* disruption on metabolic parameters in adult male mice. We show that *Atp1b4* knockout males, which survived to adulthood, have a significantly lower percentage of body fat, exhibit enhanced metabolic rate and insulin sensitivity. These robust changes in mouse metabolic parameters induced by *Atp1b4* disruption clearly demonstrate that eutherian BetaM plays an important role in the regulation of metabolism.

## RESULTS

### Construction of the BetaM knockout Mouse

To investigate the role of BetaM in muscle development *in vivo*, a BetaM-deficient mouse was generated. Gene targeted ES cells were constructed through the NIH Knockout Mouse Project (KOMP) using the knockout-first strategy based on inserting a cassette into an intron of the gene, thus resulting in a knockout at the RNA processing level (15). Germ line transmitted mice bearing the *Atp1b4* targeting knockout-first cassette inserted into intron 4 were generated by the NIH KOMP Repository at the University of California, Davis (IKMC Project Report - KOMP-CSD (ID: 23729). The *Atp1b4*-targeted ES cells were derived from C57BL/6J mice. The chimeric mice have been backcrossed to C57BL/6J mice for over 12 generations, resulting in a uniform C57BL/6J genetic background. As shown in **Fig. 1**, expression of BetaM is ablated in hemizygous males bearing the knockout-first cassette. Initial characterization of *Atp1b4* knockout mice revealed that homozygous *Atp1b4-/-* females have reproductive defects. The majority of them are sterile, and the rest have fewer pregnancies with only 3 or 4 fetuses. Heterozygous *Atp1b4-/+* females and *Atp1b4-/Y* hemizygous males are fertile and generate progeny that allow direct comparison of wild type (WT) and knockout (KO) male littermates. The survival rate of *Atp1b4-/Y* males and *Atp1b4-/-* females is 2-3 times lower (P<0.0003, Chi-square test) than that of male wild type and *Atp1b4-/+* female littermates, respectively.

**Figure 1.**
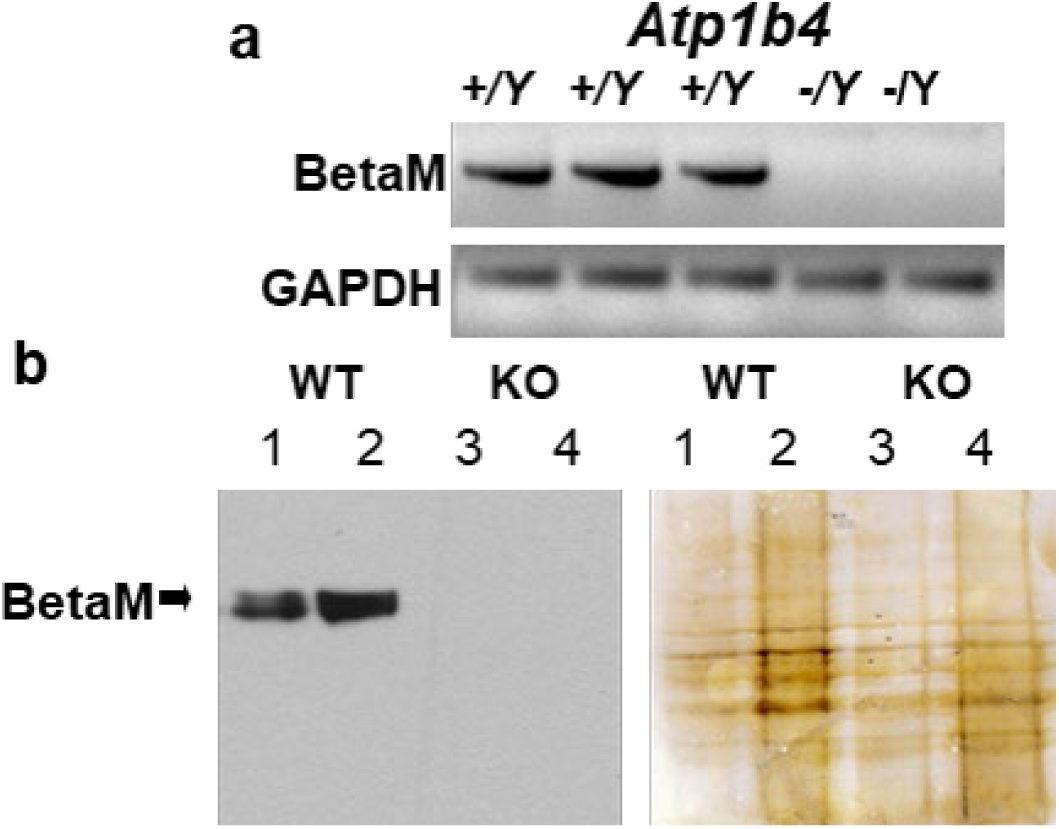
Expression of BetaM is ablated in hemizygous males bearing knockout-first cassette in intron 4. (A) RT-PCR analysis (35 cycles) of RNA from tongue skeletal muscles of 3 day old wildtype (WT) and knockout (KO littermates) using primers complementary to exons 3 and 5 of Atp1b4. GAPDH is shown as a control. (B) (Left) Immunoblot of protein from pooled tongue and hindlimb muscles of 3- day old WT and KO littermates using BetaM antibodies. (Right) Silver staining of the blot shows equal protein loading. Total protein loaded 10mgs (lanes 1 and 3) and 30mgs (lanes 2 and 4).

### Mice deficient in *Atp1b4* exhibit reduced body weight and adiposity

Birth weight has been shown to be associated with future risk of human adult metabolic disorders, including type 2 diabetes, obesity, and cardiovascular disease (16, 17). Therefore, we analyzed effects of the *Atp1b4* deletion on metabolic parameters of adult male mice. Wild-type and *Atp1b4* deficient littermates were fed a regular chow diet for 60 days starting at 2 months of age and then characterized. We first examined their body weight and visceral adiposity. As shown in **Fig. 2**, *Atp1b4* KO mice fed a regular diet (RD) have a 20% reduction in body weight and a 70% reduction is fat mass that is associated with an increase in the percent of lean mass compared with control mice. While there was a small increase in skeletal and cardiac mass to body weight, the change in body composition in KO mice was mostly due to reduction of all adipose tissue, including white adipose tissue (WAT), brown adipose tissue (BAT), visceral fat (Visc. Fat), and internal adipose tissue (int. fat.) (**Fig. 3**). These findings demonstrate that loss of *Atp1b4* dramatically reduces fat accumulation. In agreement with reduced fat mass, we also found lower sized-adiposity (data not shown), accompanied by less inflammation (data not shown) and reduced fibrosis.

**Figure 2.**
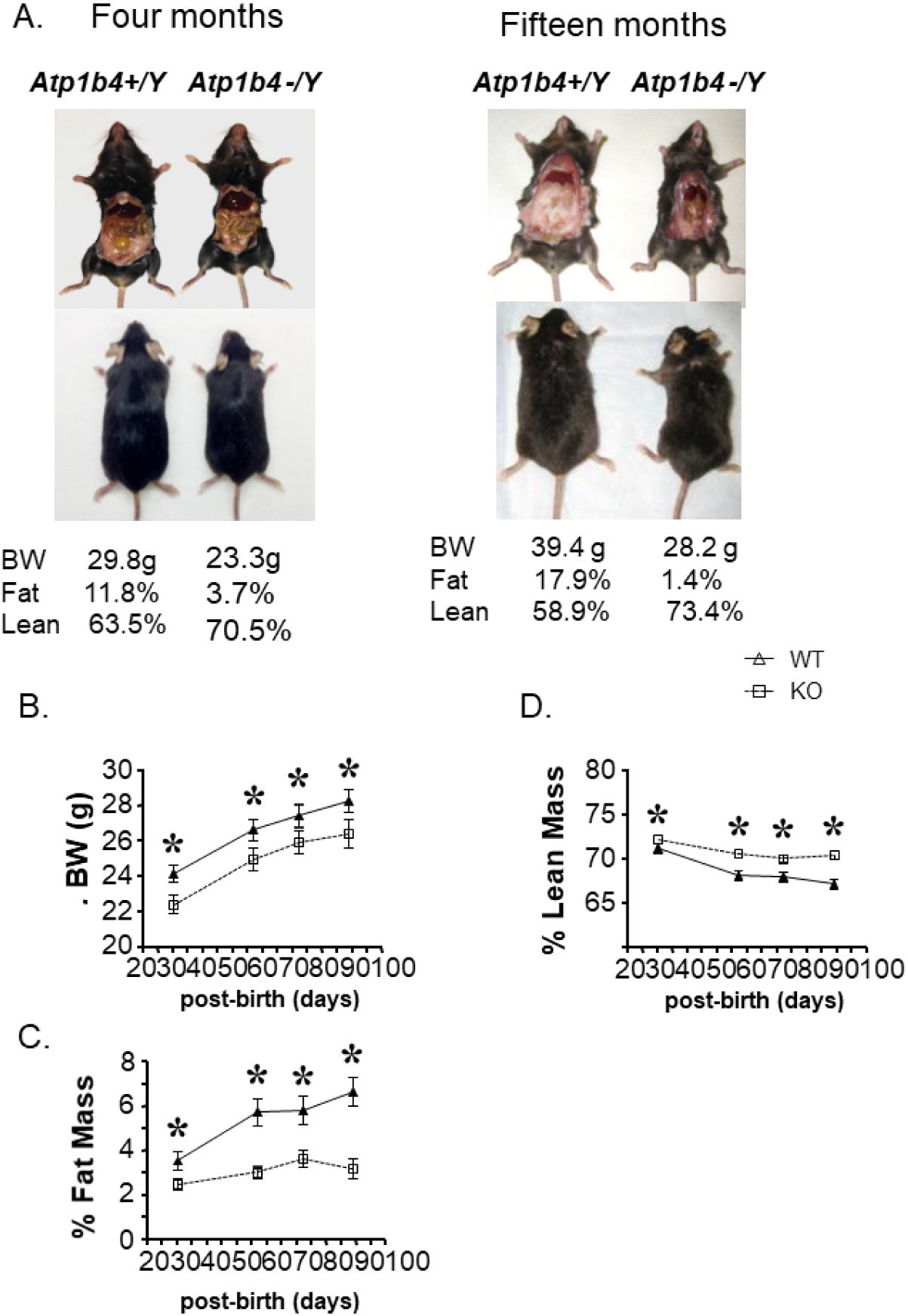
Atp1b4 KO mice have reduced body weight and altered body composition. (A) Representative pictures of 4-month old (left) and 15-month old (right) Atp1b4+/Y and Atp1b4-/Y littermates fed regular chow beginning at weaning. Four month old Atp1b4+/Y and Atp1b4-/Y mice fed RD upon weaning were assessed for: (B) Body Weight. Nuclear magnetic resonance was also performed to measure (C) % Fat Mass and (D) % Lean Mass (N=10/each group).

**Figure 3.**
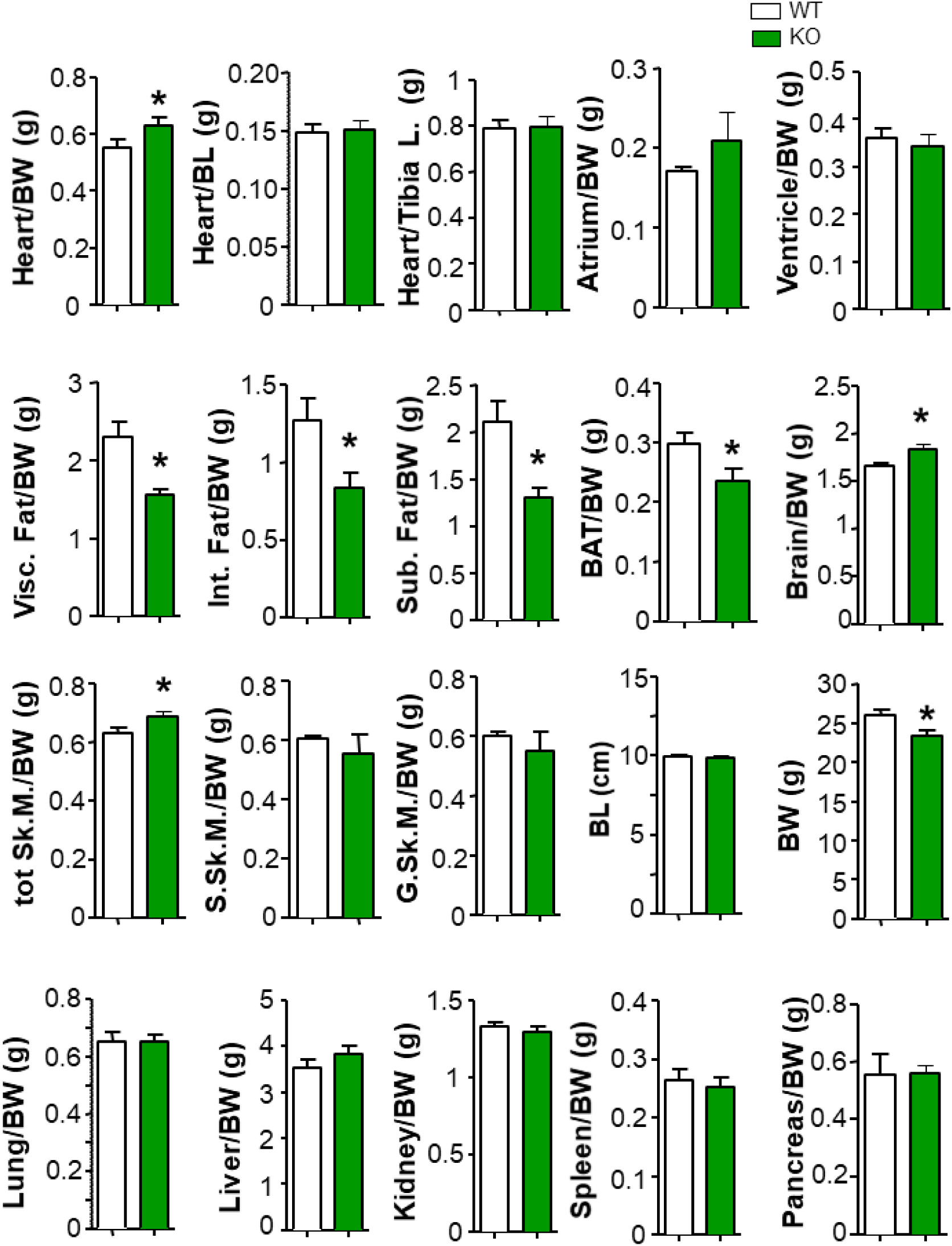
Atp1b4 KO mice have reduced adiposity. Four month old Atp1b4+/Y and Atp1b4-/Y mice fed RD upon weaning were sacrificed and the indicated tissues dissected and weighed. Tissue mass is shown relative to body weight. (N=10/each group). BW=Body weight, BL=Body length, Int. Fat=Internal fat, Visc. Fat= Visceral fat, Sub. Fat=subcutaneous fat, BAT=Brown adipose tissue. G. Sk.M=Gastrocnemius skeletal muscle, S. Sk.M=Soleus Skeletal Muscle.

### *Atp1b4* deficient mice have lower blood glucose levels, increased insulin sensitivity and glucose tolerance

Since blood glucose levels and insulin sensitivity is associated with body weight, we compared random and fasting blood levels as well as glucose tolerance and insulin sensitivity in *Atp1b4* KO mice versus WT. Atp1b4 KO had both lower random (**Table 1**) and fasting blood glucose levels after a 7 hour (**Fig. 4A, left**) or overnight fast (Fig. 4A, right). Insulin sensitivity (**Fig. 4B**) and glucose tolerance (**Fig. 4C**) also improved in *Atpb4* KO mice. We also looked at serum levels of insulin and c-peptide. Insulin levels were significantly lower in KO mice while the level of c- peptide was not significantly different between WT and KO mice. Glucose levels measured at fasting were lower than wild-type mice, both upon RD or HF diet (**Table 1**). Both fasting and random blood glucose levels were found to be reduced in *Atp1b4* knock-out mice compared with controls (**Table 1**). Taken together, these data demonstrate that deficiency in *Atp1b4* improves glucose metabolism and insulin action.

**Figure 4.**
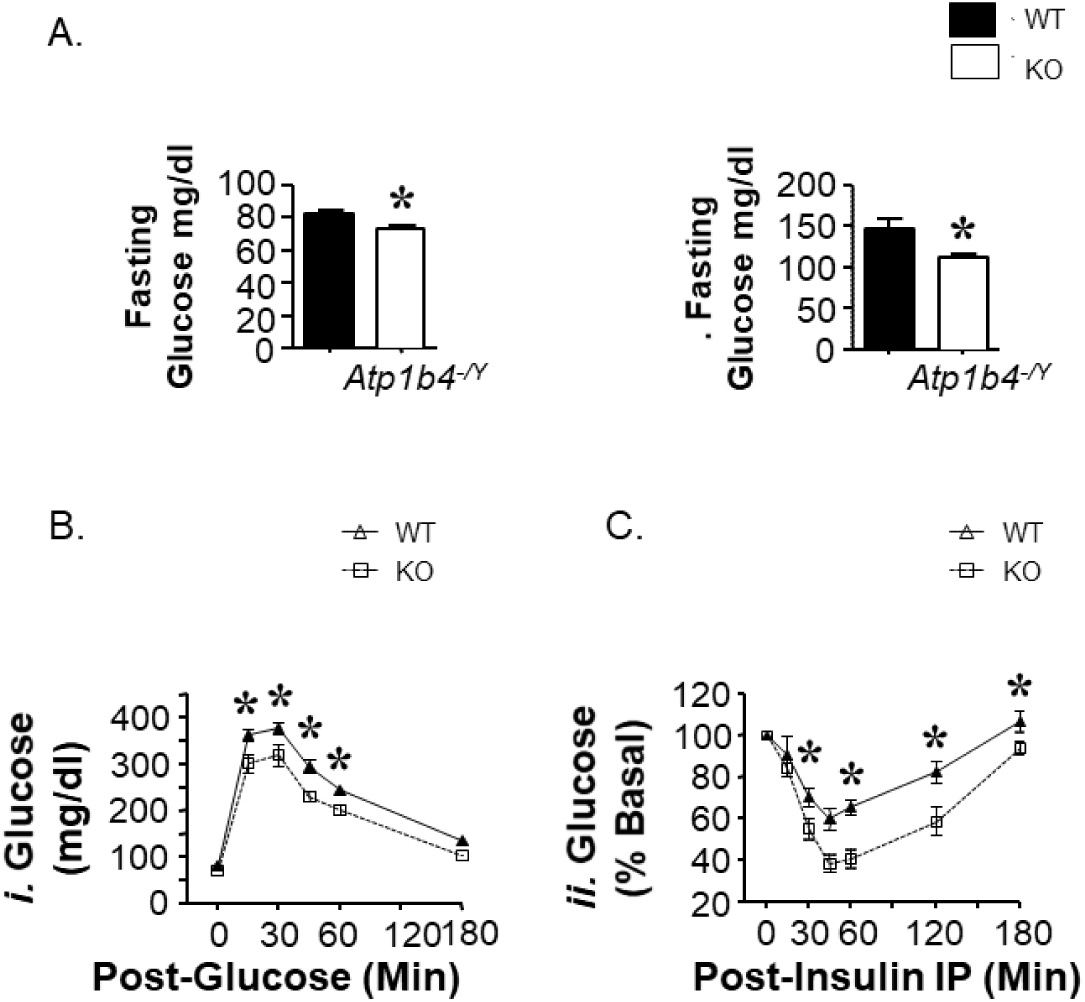
Atp1b4 KO mice have lower fasting blood glucose levels, improved insulin sensitivity and glucose tolerance. (A) Blood glucose levels in 4 month old WT and Atp1b4 KO mice were measured after an overnight (left) or a 7 hour fast (right). (B) Blood glucose levels were measured after fasted WT or KO mice received intraperitoneal injections of glucose (1.5g/kg−1) for 0-120 minutes and their tail vein blood glucose measured. (N=11/each group) (C) Blood glucose levels were measured after fasted mice received intraperitoneal injections of insulin (0.75 U kg−1). Measurements are presented as percent of basal values. (N=7 WT and 9 KO). Values are expressed as means ± SEM. *P≤0.05 (N=11/each group).

**Table 1.** Effect of Atp1b4 deficiency on metabolic parameters.

| **Effect of *Atp1b4* deficiency on metabolic parameters** |  |  |
| --- | --- | --- |
|  | **<u>*Atp1b4*<sup>+/Y</sup></u>** | **<u>*Atp1b4*<sup>-/Y</sup></u>** |
| **Fasting plasma** |  |  |
| **Insulin (pmol/L)** | 38.82 ± 1.69 | 34.30 ± 0.92* |
| **C-peptide (pmol/L)** | 520.8 ± 73.59 | 525.9 ± 79.53 |
| **C-peptide-to-insulin molar ratio** | 16.02 ± 2.22 | 15.73 ± 2.44 |
| **Blood glucose (mg/dL)** |  |  |
| **Fasting** | 81.86 ± 2.03 | 72.73 ± 2.05** |
| **Random** | 166.3 ± 6.10 | 144.5 ± 5.16* |

**Table 2.**
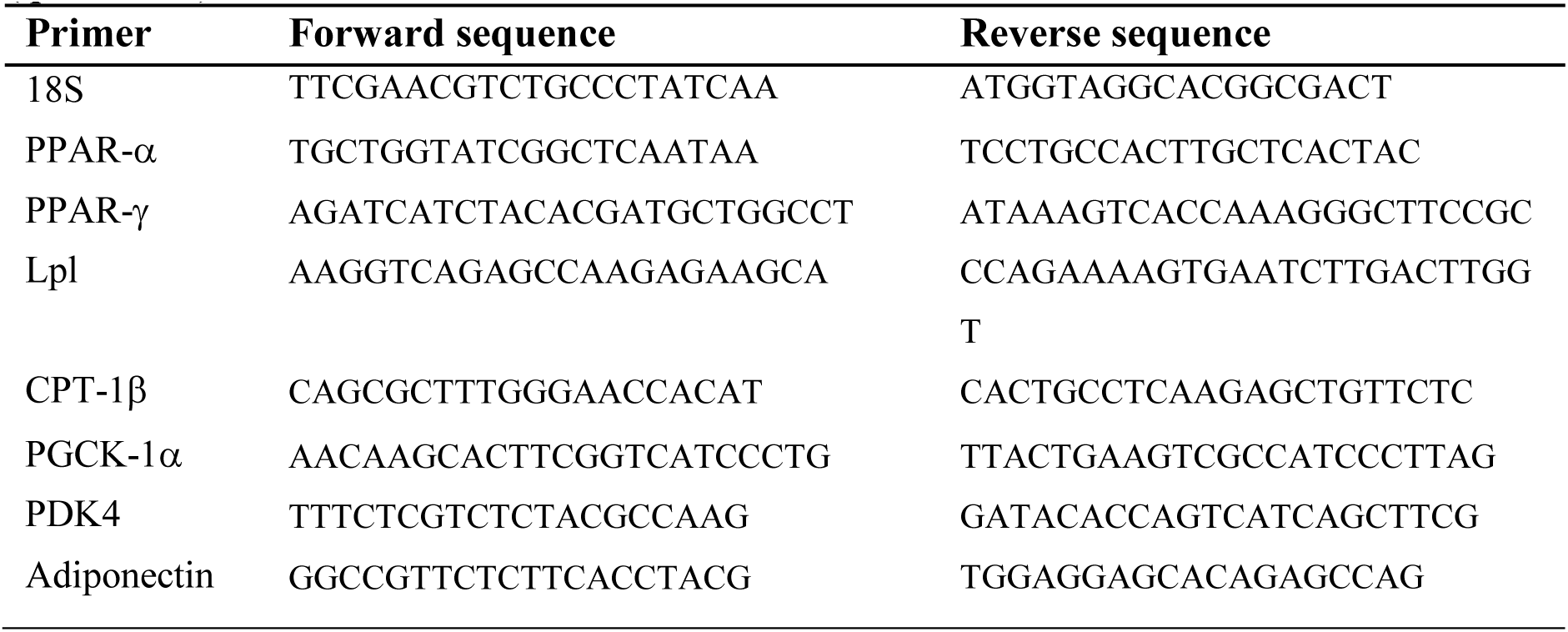
Primer sequences used for semi-quantitative Real Time PCR Polymerase Chain Reaction (qRT-PCR)

### Altered substrate oxidation and gene markers for β-oxidation and lipid accumulation in *Atp1b4* deficient mice

Next, we wanted to determine whether *Atp1b4* KO mice have elevated thermogenesis, lipid oxidation, or activity. To this end, by measuring fatty acid oxidation in skeletal muscle we found increased energy metabolism in *Atp1b4* KO mice compared with WT controls (**Fig. 5A**). To confirm increased lipid oxidation, we also measured gene expression markers for β-oxidation and lipid accumulation. We found that *Atp1b4* knock-out mice showed increased mRNA levels for β- oxidation in muscle (**Fig. 5B**). Peroxisome proliferator-activated receptor alpha (PPARα) is a major regulator of lipid metabolism, activation of PPARα promotes uptake, utilization, and catabolism of fatty acids by upregulation of genes involved in fatty acid transport, fatty acid binding and activation, and peroxisomal and mitochondrial fatty acid β-oxidation (18). Some of the genes regulated by PPARα include Pyruvate Dehydrogenase Kinase 4 (PDK4), PDK4 inhibits the pyruvate dehydrogenase complex by phosphorylating one of its subunits contributing to the regulation of glucose metabolism (19), and Carnitine Palmitoyltransferase 1B (CPT-1b), CPT-1b is required for the net transport of long-chain fatty acyl-CoAs from the cytoplasm into the mitochondria (20), both are upregulated, 2.5 and 2 fold respectively, in the muscle of the *Atp1b4* knock-out mice. In addition, *Atp1b4* knock-out mice exhibit lower expression of mRNA levels for lipid accumulation (Fig.5b), such as Peroxisome proliferator-activated receptor gamma (PPARγ). PPARγ regulates fatty acid storage and glucose metabolism (21). When activated, PPARγ stimulates lipid uptake and adipogenesis by fat cells and upregulation of adipogenic genes. Lipoprotein Lipase (Lpl), Lpl functions as a homodimer, and has the dual functions of triglyceride hydrolase and ligand/bridging factor for receptor-mediated lipoprotein uptake (22), and adiponectin, adiponectin circulates in the plasma and is involved in metabolic processes including glucose regulation and fatty acid oxidation (23), are both downregulated in *Atp1b4* KO mice.

**Figure 5.**
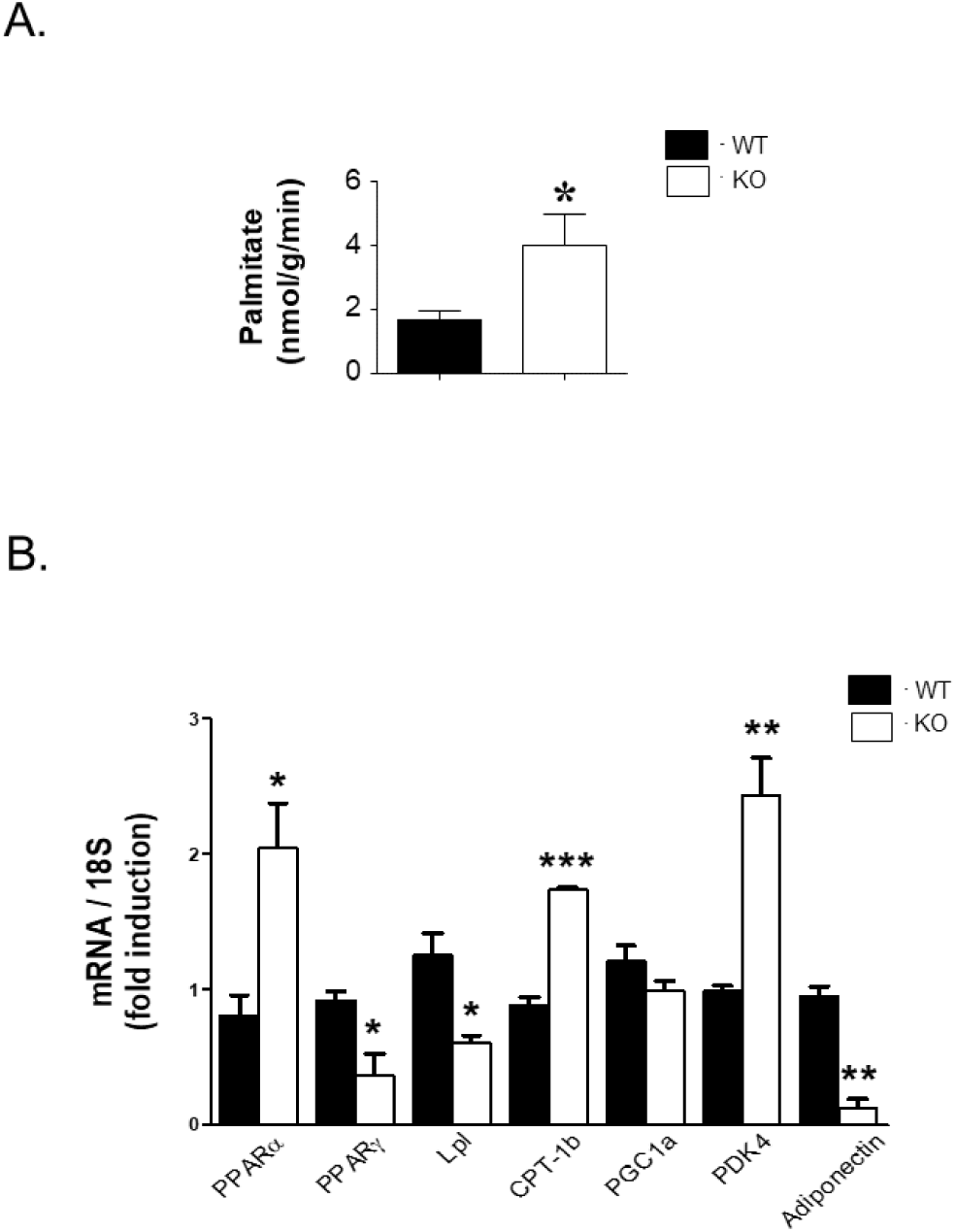
Fatty acid oxidation increases in muscle from Atp1b4 KO mice. (A) Fatty acid oxidation in skeletal muscle in 4 month-old male Atp1b4+/Y and Atp1b4-/Y mice fed RD (N=6/each group). (B) Analysis of skeletal muscle from Atp1b4+/Y and Atp1b4-/Y mice for expression of gene markers involved in thermogenesis and lipid oxidation. Transcript expression was normalized to 18S RNA. Values are expressed as means ± SEM. *P≤0.05, **P< 0.01, ***P<0.005 versus Atp1b4+/Y.

### Elevated energy expenditure in *Atp1b4* deficient mice

To confirm the results of increased energy metabolism in *Atp1b4* KO mice, the metabolic rate in these mice was evaluated. Consistently, *Atp1b4* KO mice maintained higher energy expenditure (**Fig. 6A**), higher O2 consumption (**Fig. 6B**), higher CO2 production (**Fig. 6C**). Moreover, KO mice had lower respiratory exchange ratios, suggesting a greater use of fat as an energy source (**Fig. 6D**). KO mice showed increased spontaneous locomotor activity (**Fig. 7A**) and increased food consumption (**Fig. 7B**) compared with the control counterparts.

**Figure 6.**
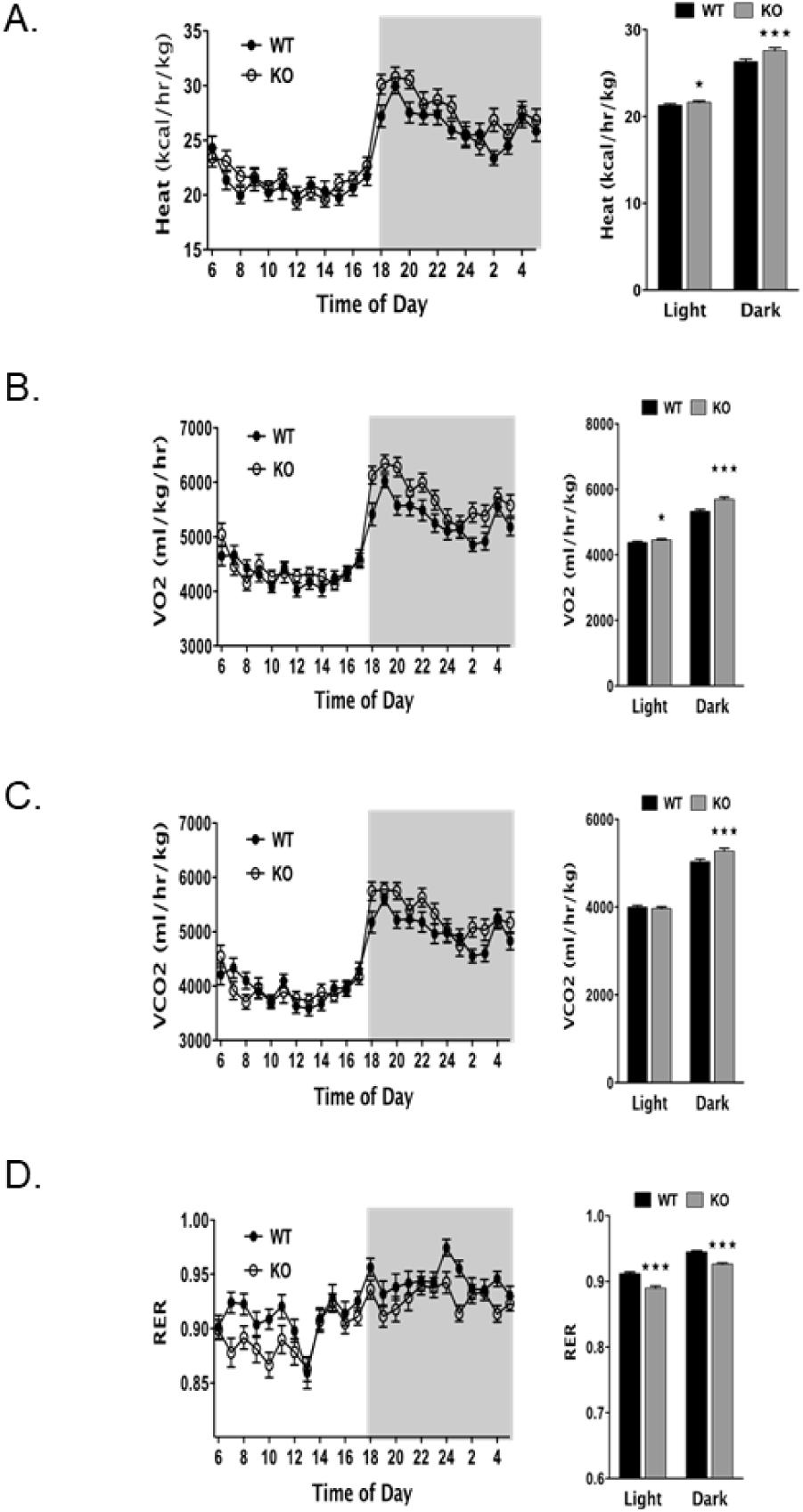
Increased energy metabolism in Atp1b4 KO mice. 3 month-old male Atp1b4+/Y and Atp1b4-/Y mice fed a RD diet were individually caged (N=8/each group), given free access to food, and subjected to indirect calorimetry analysis in a 24-h period for 5 days to assess (A) heat generation or energy expenditure [EE] (kcal/h/kg lean mass) (B) VO2 consumption (ml/h/kg lean mass), (C) VCO2 production (ml/h/kg lean mass) (D) respiratory exchange ratio, RER. Values are expressed as means ± SEM. *P≤0.05, **P< 0.01, ***P<0.005 versus Atp1b4+/Y.

**Figure 7.**
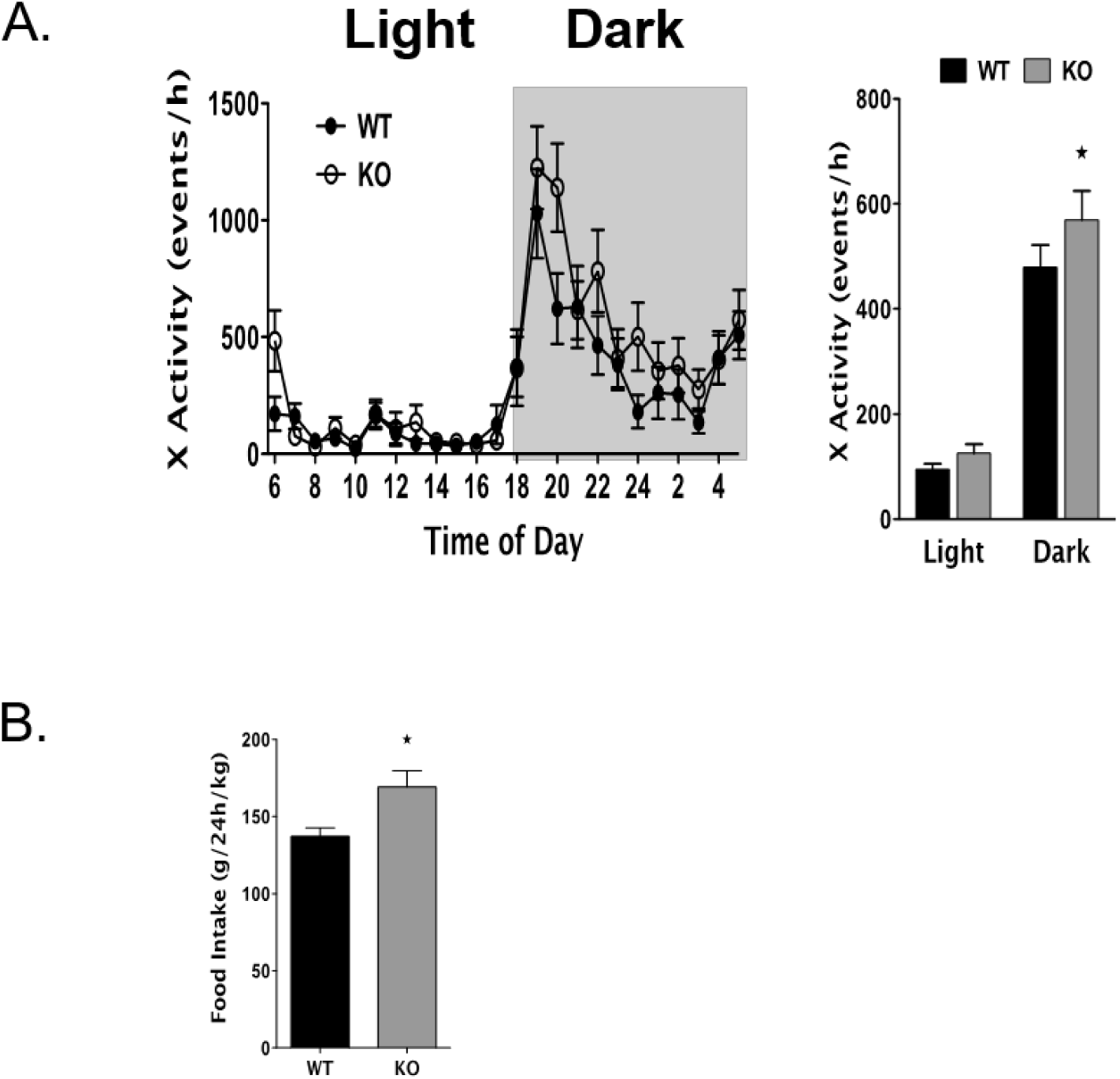
3 month old Atp1b4 KO male mice are more active and are hyperphagic. (A) Spontaneous activity. (B) Food intake Values are expressed as means ± SEM. *P≤0.05, **P< 0.01, ***P<0.005 versus Atp1b4+/Y.

## DISCUSSION

The *ATP1B4* genes, members of the X,K-ATPase β-subunit gene family, are a rare example of orthologous vertebrate gene co-option (2). This evolutionary phenomenon has led to profound alterations in the functional properties, significantly transforming the structure proteins and expression pattern of encoded BetaM (1, 5–7, 10–12). Expression of eutherian BetaM is tissue-specific, the highest level is in skeletal muscle, a lower level in heart and a much lower level in skin (5–7, 10). In muscle, BetaM expression is temporally regulated; it is robustly induced in the last quarter of pregnancy, remains high in neonatal muscle, and is undetectable in adult mouse muscle. This pattern of expression suggests that the function of BetaM is linked to myogenic developmental processes (5–7, 10, 11, 24–26). We previously showed that BetaM interacts with the nuclear transcriptional co-regulator Ski-interacting protein (SKIP) in neonatal muscle, influencing the activity of TGF-β-responsive reporters. BetaM binds to the promoter of Smad7, a critical negative regulator of TGF-β/Smad signaling and up-regulates its expression (1). These findings provided the first experimental evidence that eutherian BetaM functions as a regulator of transcription. We recently reported that BetaM binds to the MyoD promoter *in vivo* and found that exogenously expressed BetaM enhanced MyoD expression in cultured myoblasts, indicating a regulatory role in myogenesis (14). BetaM promoted epigenetic changes and recruited the SWI/SNF chromatin remodeling subunit, BRG1. These results indicate that eutherian BetaM regulates muscle gene expression by promoting changes in chromatin structure.

To gain an understanding of the function of BetaM *in vivo*, we generated a whole body BetaM KO mouse. Surprisingly, the absence of BetaM in neonatal hemizygous male mice significantly affected whole-body metabolism. Because BetaM expression is restricted to muscle in the neonatal muscle, this suggests that the loss of BetaM in muscle impacts whole body metabolism. Skeletal muscle, a major site for the regulation of fatty acid and glucose metabolism, plays a significant role in the control of obesity, diabetes, and cardiovascular disease, and is itself regulated by whole body metabolism (27–29). As the largest organ, skeletal muscle is responsible for approximately 80% of glucose uptake from the circulation. Obesity and a sedentary lifestyle lead to lipid accumulation and metabolic alterations in skeletal muscle, which impede glucose regulation and contribute to the development of insulin resistance (30, 31). Skeletal muscle also releases myokines, which mediate tissue crosstalk and influence whole-body metabolism. These myokines can either increase or decrease inflammation, and insulin resistance (15, 16). However, the impact of muscle on whole-body metabolism have not been extensively investigated and the mechanisms remain to be elucidated.

While our previous studies have indicated that BetaM expression is temporally expressed, we did see activation of BetaM during muscle regeneration. A recent study also detected activation of BetaM expression in incisional hernia tissue, concommittantly with non-alcoholic fatty liver disease-related, TNF, and IL-17 signaling pathways (32). Collectively, these studies suggest that BetaM expression is induced in adults and may influence muscle metabolism, potentially affecting whole-body metabolism. Currently, we are investigating the mechanisms that regulate BetaM expression and its impact on overall metabolism in adults.

Plasma insulin, C-peptide, C-Peptide-to-Insulin Molar Ratio, and glycemia levels were measured in 4 month-old male Atp1b4+/Y and Atp1b4-/Y fed RD for 2 months. Mice were subjected to an over-night fasting before being sacrificed (N=10/each group). Values are expressed as means ± SEM. *P≤0.05, **P< 0.01, ***P<0.005 versus Atp1b4+/Y..

## MATERIALS AND METHODS

### *Atp1b4* knock-out mouse model

Gene targeted ES cells have been constructed through the framework of the NIH Knockout Mouse Project (KOMP) using the knockout-first strategy based on inserting a cassette into intron of the gene that produces a knockout at the RNA processing level (15). Germ line transmitted mice (C57BL/6N) bearing *Atp1b4* targeting knockout-first cassette were generated by contract with the NIH KOMP Repository at the University of California, Davis (IKMC Project Report - KOMP-CSD (ID: CSD23729). The *Atp1b4* targeted ES cells were from C57BL/6J mice. The chimeric mice ere bred to C57BL/6J mice for more than 12 generations resulting in mice with a uniform C57BL/6J genetic background. Mice were housed under a 12-12 hour light-dark cycle (6 am light on) with *ad libitum* access to tap water, a standard chow (Catalog #2016S, 12 kcal% fat, Harlan Teklad Laboratory Diets, WI, USA). The Institutional Animal Care and Use Committee (IACUC) approved all experimental protocols.

### Body Composition

Whole body composition was assessed by Nuclear Magnetic Resonance technology (Bruker Minispec; Billerica, MA, USA).

### Intra-peritoneal Glucose Tolerance Test

Mice were fasted overnight from 5 pm to 11 am. Fasted blood glucose was taken from the tail vein by snipping the tail. Glucose at 1.5 g/kg body wt (50 % dextrose solution) was administered by intra-peritoneal (IP) injection to conscious animals. Blood glucose was measured from the tail vein at 15, 30, 45, 60 and 120 minutes post-glucose injection. Glucose levels are expressed as mg/dl.

### Intra-peritoneal Insulin Tolerance Test

Mice were fasted for 7 hours starting at 7 a.m. Fasted blood glucose was taken from the tail vein by snipping the tail. Awake mice were IP injected with 0.75 U/kg body weight of Human insulin Novolin (Novo Nordisk, Princeton, NI, USA; NDC 0169-1833-11). Blood glucose was measured from the tail vein at 0, 15, 30, 45, 60, 120 and 180 minutes post insulin injection. Glucose levels are expressed as percentage to fasting levels.

### Metabolic parameters

Eye puncture by retro-orbital bleeding was applied in order to withdraw blood at 1100h from overnight fasted mice. Blood was collected into heparinized micro-hematocrit capillary tubes (Fisherbrand 22-362-566) and immediately chilled on ice. After 30 min of centrifugation at 3000 rpm and 4°C, plasma was stored at −80°C. Plasma insulin and C-peptide were measured by using the corresponding mouse radio-immunoassay kit (Linco Research, St Charles, MO, USA). Intra- and inter-assay coefficient of variation for insulin were 2.7-5.8% and 4.0-11.2% respectively.

### Quantitative Real Time-PCR (qRT-PCR)

Mice were sacrificed and hindleg muscle tissue was collected, snap-frozen in liquid nitrogen and stored at −80°C for measurement of mRNA content by real time qPCR (ABI StepOnePlus Real-Time PCR System, Applied Biosystems, Foster City, CA). Total RNA was extracted from frozen tissue samples using PerfectPure RNA Tissue Kit–50 (5 PRIME −2900304; 2900317- Inc., MD, USA) and its quantity and purity were determined by absorbance at 260 and 280 nm. cDNA templates for qRT-PCR were synthesized using 1 μg of total RNA, 10X DNase Reaction Buffer, DNase 1 Amp Grade, dNTPs, Random Primers, and Superscript III (Invitrogen). qRT-PCR was performed using SYBR Green Master Mix (Smart Bioscience, Maumee, OH USA). All primers were used at a final concentration of 10 mΜ (Table 3). Expression level of each experimental gene was normalized to the concentration of the house-keeping gene 18S in each sample.

### Ex vivo palmitate oxidation

Skeletal muscle of overnight-fasted mice was processed as previously described (33, 34) with some modifications. Fatty acid synthase (FAS) activity was measured immediately in the presence of 0.1 µCi [14C] malonyl coenzyme A (CoA; Perkin Elmer, Waltham, MA), 25 nmol malonyl CoA, and 500 µmol/L reduced nicotinamide adenine dinucleotide phosphate. FAS activity was calculated as cpm of [14C] incorporated/mg of lysates.

### Energy balance measurements

Mice were placed in a customized indirect calorimetry system (CLAMS system, Columbus Instrument, Columbus, OH) and energy expenditure (EE), oxygen consumption (VO_2_), CO_2_ production (CO_2_), spontaneous physical activity (SPA), respiratory exchange ratio (RER) and food intake (FI) were monitored simultaneously on line for 3 consecutive days after 2 days adaptation. Measurements were taken every 20 minutes at flow rate of 0.5 l/min.

### Statistical Analysis

Quantitative data are presented as mean ± standard error of the mean (S.E.M). Values were analyzed for statistically significant differences using one-way ANOVAs and post-hoc Tukey test or two-tailed unpaired t-test. *P<0.05* was considered statistically significant (Graph Pad Prism software 5.0).

## Acknowledgements

We are grateful to Drs Lance A. Stechschulte, Larissa V. Fedorova, Nikolay B. Pestov and Nisar Ahmad for the help in initial characterization of Atp1b4 knout mouse model and to Drs. David J. Kennedy and Steven T. Haller for helpful discussions and valuable comments on manuscript. Our special thanks go to Jennifer Kalisz and Melissa Kopfman for excellent technical assistance.

The mouse strain (ID: CSD23729) used for this research project was generated from ES Cells obtained from the KOMP Repository, a NCRR-NIH supported strain repository. ES cells were created by the CSD consortium from funds provided by the trans-NIH Knock-Out Mouse Project (KOMP). To inquire about KOMP products go to www.komp.org

All authors have read and agreed to the published version of the manuscript.

## Institutional Review Board Statement

All animal experiments were conducted in accordance with the National Institutes of Health (NIH) Guide for the Care and Use of Laboratory Animals approved by The University of Toledo Institutional Animal Care and Use Committee.

